# Stratification of Risk of Progression to Colectomy in Ulcerative Colitis using Measured and Predicted Gene Expression

**DOI:** 10.1101/2021.04.02.438187

**Authors:** Angela Mo, Sini Nagpal, Kyle Gettler, Talin Haritunians, Mamta Giri, Yael Haberman, Rebekah Karns, Jarod Prince, Dalia Arafat, Nai-Yun Hsu, Ling-Shiang Chuang, Carmen Argmann, Andrew Kasarskis, Mayte Suarez-Farinas, Nathan Gotman, Emebet Mengesha, Suresh Venkateswaran, Paul A. Rufo, Susan S. Baker, Cary G. Sauer, James Markowitz, Marian D. Pfefferkorn, Joel R. Rosh, Brendan M. Boyle, David R. Mack, Robert N. Baldassano, Sapana Shah, Neal S. LeLeiko, Melvin B. Heyman, Anne M. Griffiths, Ashish S. Patel, Joshua D. Noe, Sonia Davis Thomas, Bruce J. Aronow, Thomas D. Walters, Dermot P. B. McGovern, Jeffrey S. Hyams, Subra Kugathasan, Judy H. Cho, Lee A. Denson, Greg Gibson

## Abstract

An important goal of clinical genomics is to be able to estimate the risk of adverse disease outcomes. Between 5% and 10% of ulcerative colitis (UC) patients require colectomy within five years of diagnosis, but polygenic risk scores (PRS) utilizing findings from GWAS are unable to provide meaningful prediction of this adverse status. By contrast, in Crohn’s disease, gene expression profiling of GWAS-significant genes does provide some stratification of risk of progression to complicated disease in the form of a Transcriptional Risk Score (TRS). Here we demonstrate that both measured (TRS) and polygenic predicted gene expression (PPTRS) identify UC patients at 5-fold elevated risk of colectomy with data from the PROTECT clinical trial and UK Biobank population cohort studies, independently replicated in an NIDDK-IBDGC dataset. Prediction of gene expression from relatively small transcriptome datasets can thus be used in conjunction with transcriptome-wide association studies to stratify risk of disease complications.

## INTRODUCTION

Genetic risk assessment in humans has to date focused mainly on prediction of disease onset (1), whereas arguably the greater clinical need is for prediction of disease progression (2,3). Polygenic risk scores (PRS) may sometimes meet both needs, such as the ability of a PRS for coronary artery disease to stratify people with respect to the likely effectiveness of statins or PCSK9 inhibitors (4–6). This is not generally expected to be the case, however, and in the context of inflammatory bowel disease, there appears to be little influence of the heritability for disease on progression to complicated disease (7). Since genome-wide association studies sufficiently powered to develop accurate PRS for progression or therapeutic response are not yet available, there is a need for alternative genomic strategies.

A promising approach is gene expression profiling, which very often discriminates cases and controls. For both Crohn’s disease and ulcerative colitis, RNAseq of ileal and rectal biopsies respectively, generates discriminators of disease severity and progression to complications or remission that are at least as good as clinical indices (8–10). Combining eQTL with GWAS signals with RNAseq data also supports transcriptional risk scores (TRS), namely weighted sums of polarized z-scores of transcript abundance, that predict stricturing or penetrating Crohn’s disease (11). As profiling moves to the single cell level, it is clear that gene expression will also define the identities of critical cell types in which pathogenic alleles act (12–14) and likely refine transcript-based risk assessment. The main limitation of this approach is the ability to obtain appropriate tissue biopsies.

Consequently, transcriptome-wide association studies (TWAS) have been proposed to fill this gap (15,16). These are analyses that essentially sum the cis-eQTL effects at a locus in order to predict gene expression in a case-control cohort where only genotypes are available. Differential expression predictions have been shown to highlight candidate genes for a range of disease (17). Here we demonstrate that the further utility of TWAS to generate a predicted polygenic transcriptional risk score (PP-TRS) for ulcerative colitis, which not only discriminates cases, but also progression to major disease complication requiring colectomy for up to 10% of patients (18–20). Genomic analysis of just hundreds of individuals, projected onto the UK Biobank (21), supports polygenic risk assessment that outperforms the current PRS for ulcerative colitis. Our analyses also provide insight into the cell-type specificity in both epithelial and immune compartments for IBD-GWAS loci.

## RESULTS AND DISCUSSION

PROTECT is a multicenter pediatric inception cohort study of response to standardized colitis therapy^**9a**^. We have previously shown that a signature of rectal mucosal gene expression at diagnosis, prior to therapeutic intervention, associates with corticosteroid-free remission with mesalamine alone observed in 38% of 400 patients by week 52 of follow-up^**9**^. A signature of rectal mucosal gene expression associated with week 4 corticosteroid response in PROTECT is related to one indicative of response to anti-TNFα and anti-α_4_β_7_ integrin therapy in adults^**10**^, and reciprocally, active pediatric UC was associated with suppression of mitochondrial gene expression, and increasing disease severity with elevated innate immune function. In order to more explicitly model progression to colectomy observed in 6% (25 of 400) of the patients within one year of diagnosis, we performed differential expression analysis between baseline rectal RNAseq biopsies of 21 patients who progressed to colectomy, and 310 who did not. The volcano plot in Fig. 1a shows down-regulation of 783 transcripts in the colectomy cases (red), and up-regulation of 1,405 transcripts (blue) at the experiment-wide threshold of p<4×10^−6^. Gene set enrichment analysis^**22**^ summarized in Fig. 1b highlights engagement of multiple pathways previously implicated in adverse outcomes in inflammatory bowel disease, including TNF and interferon signaling, and various signatures of inflammation and immune response^**8,23**^.

**Figure 1.**
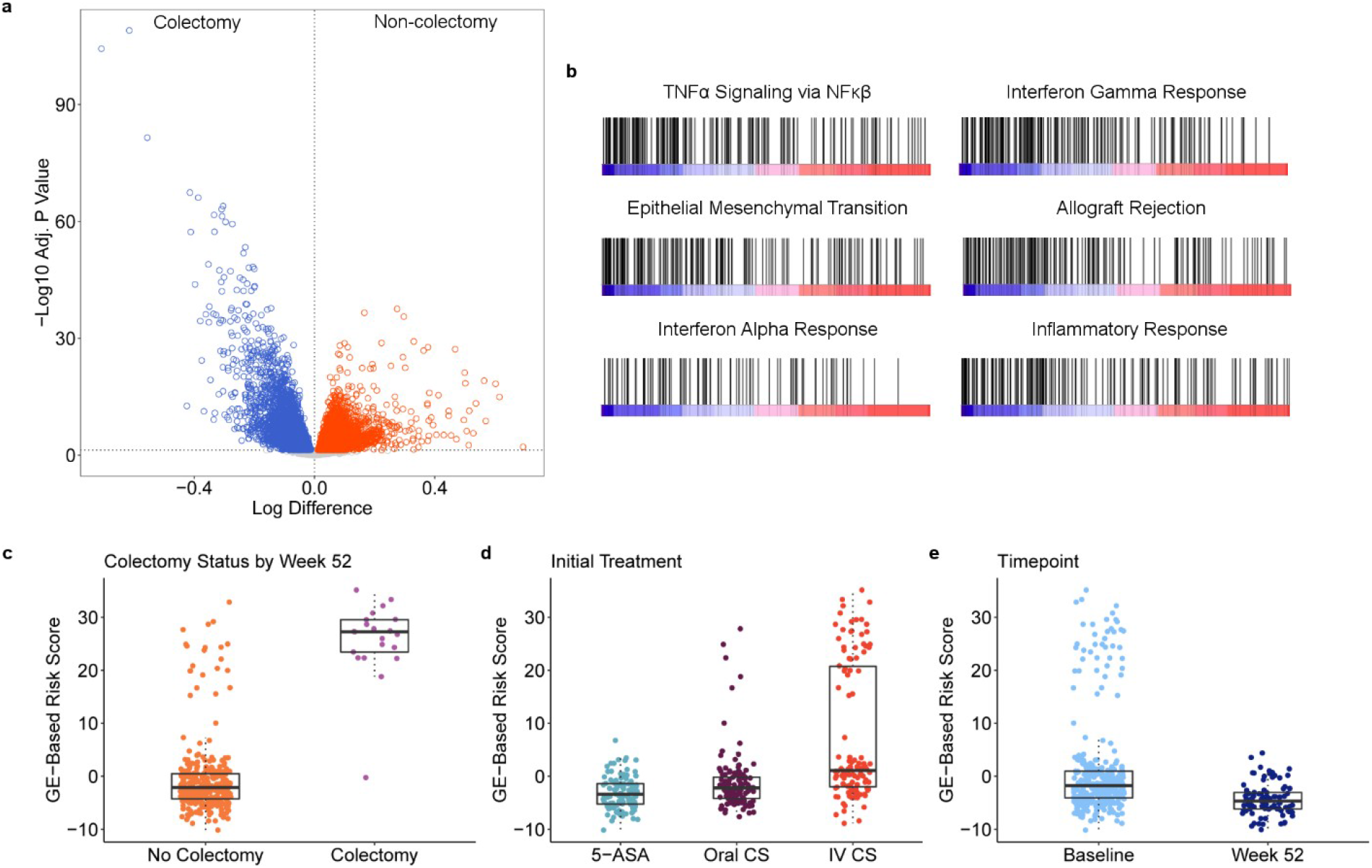
Differential Expression Associated with Colectomy in the PROTECT study. (a) Volcano plot of significance (negative log10 of the p-value) against difference in expression on log2 scale, with genes up-regulated in colectomy in blue. (b) Six pathways highlighted by gene set enrichment analysis as up-regulated in colectomy. Each bar represents a gene in the indicated pathway, and position along the axis is representative of rank order of differential expression. From left to right, top to bottom, FDR < 10^−4^, < 10^−4^, < 10^−4^, < 10^−4^, 2.4×10^−4^ and 2.0×10^−4^. A full list of pathways can be found in Table S2. PC1 of the differentially expressed genes as a function of (c) colectomy status at week 52; p = 2×10^−45^, (d) initial treatment; p = 5×10^−20^, and (e) baseline or week 52 follow-up biopsy profile; p = 2×10^−7^. All boxplots indicate 1^st^ and 3^rd^ quartile as box ends, with center median line and whiskers extending to farthest point within 1.5 times the interquartile range.

The first principal component (PC1_col_) of the top 150 of these differentially expressed genes has a weak negative correlation with our previously reported signature of remission detected in a subset of 206 patients using a different RNAseq protocol^**10**^. With very high significance, it distinguishes the colectomy cases from non-progressors, as all but one case have PC1 scores greater than 10, a value exceeded by only 20 of the 317 non-colectomy cases (Fig. 1c). This PC1_col_ predictor is orders of magnitude more significant than observed with similar scores derived by 1000 permutations of the data (Fig. S1). All of the high PC1_col_ individuals were placed initially on corticosteroids, the majority intravenously (Fig. 1d); the score also correlates with a gradient of disease severity indicated by baseline PUCAI (pediatric ulcerative colitis activity index)^**24**^ and initial treatment. We also obtained rectal biopsy RNAseq data for 92 patients at week 52 and observed significant depression of the score (Fig. 1e), indicative of mucosal healing even in the cases with elevated initial gene activity (none of the follow-up cases were colectomy, since the surgical procedure had been performed earlier than week 52). Figure S2 shows that PC1 remains associated with Mayo endoscopic score (25) even at week 52, and that the change in PC1 molecular score over time correlates with the degree of mucosal healing.

Given the marked shift in gene expression at follow-up, we next asked whether local regulation of the gene expression might contribute, by performing comparative eQTL analysis. Figure 2a indicates generally high concordance in the effect sizes (betas) at both time-points, with slight inflation of the estimates at baseline (1,416 blue effects) or week 52 (421 magenta effects), likely due to winner’s curse. There were 72 eSNPs significantly regulating 308 genes at both time points, with the smaller number of eQTL at week 52 attributable to the smaller sample size. One quarter of the baseline eQTL are at least 2-fold greater than at week 52, and one third of the follow-up eQTL are at least 2-fold greater than at baseline. Clearly visible in Fig 2a are 33 apparently week 52-specific effects that are more than 20-fold greater than at baseline, the majority with reduced expression of the minor allele. Examples of baseline and follow-up specific eQTL affecting a variety of gene functions in immunity and epithelial cell biology are shown in Fig. 2b. Some of the change in eQTL profiles is likely attributable to an increase in the proportion of epithelial relative to immune cells at week 52 (Fig. S3).

**Figure 2.**
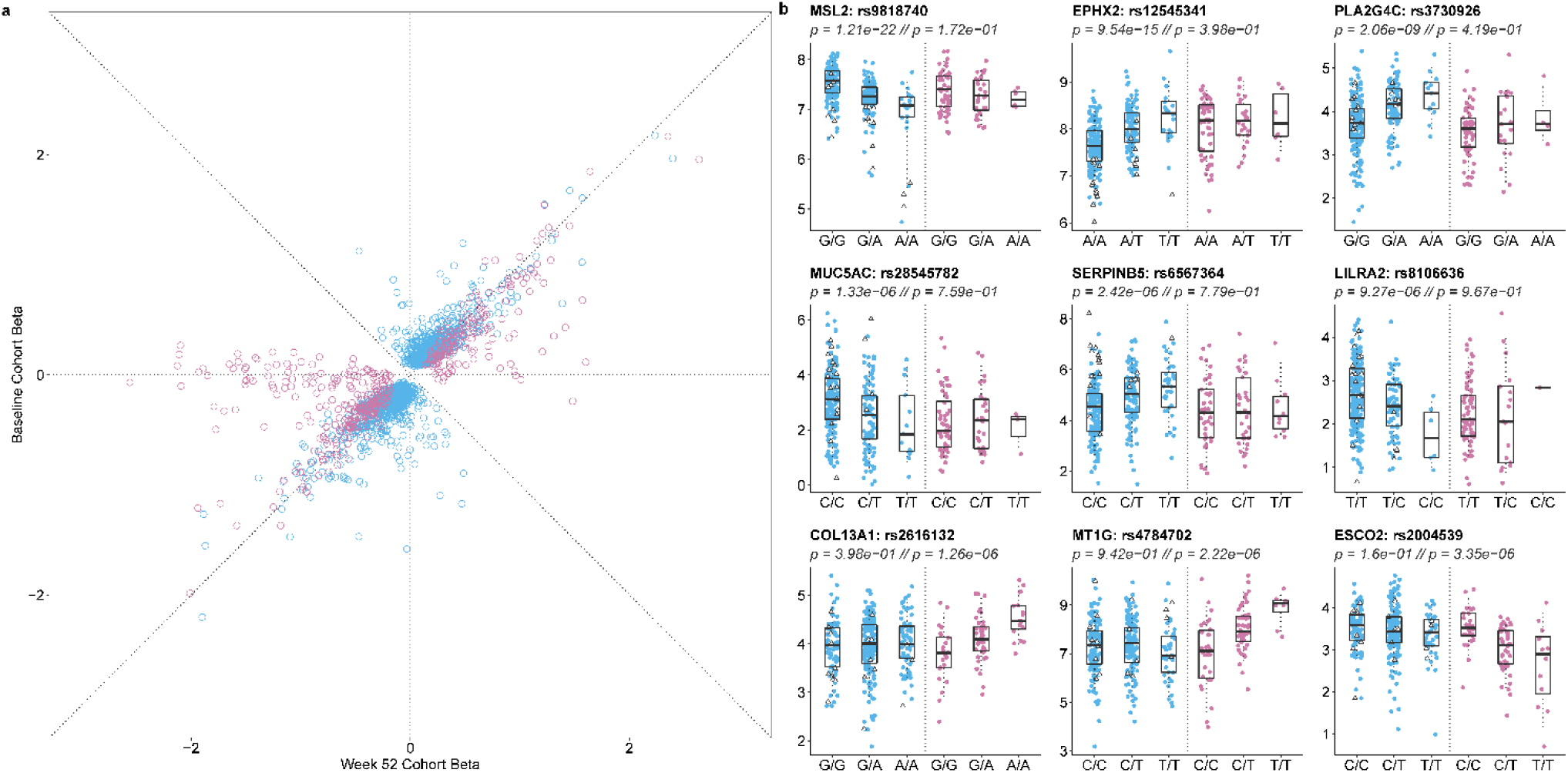
eQTL contrast between baseline and week 52 follow-up in the PROTECT study. (a) Comparison of effect sizes (betas) for the effect of the minor allele on gene expression. Blue eQTL were discovered at baseline, and magenta only at week 52. (b) Examples of nine genes with differential eQTL effects at the two timepoints showing observed transcript abundance as a function of genotype at baseline or week 52 follow-up. The bottom row are genes with eQTL only at follow-up. All boxplots indicate 1^st^ and 3^rd^ quartile as box ends, with center median line and whiskers extending to farthest point within 1.5 times the interquartile range. Note that many of the genes with large negative follow-up betas in panel (a) have relatively small minor allele frequencies, hence insufficient homozygous minor allele genotypes to plot. A full list of peak eQTL can be found in Table S3.

Next, we asked whether the intersection of GWAS, eQTL and differential expression could be used to generate a transcriptional risk score (TRS) for colectomy, analogous to the one we recently developed for prediction of risk of progression to complicated Crohn’s disease^**11**^. The heatmap in Fig. 3a showing the abundance of 26 transcripts included in the TRS_IBD_ derived with *coloc* overlap (26) of IBD GWAS and peripheral blood eQTL signals, indicates striking enrichment for elevated or reduced expression of a dozen transcripts in the baseline rectal biopsies of PROTECT patients destined for colectomy. The strongest clusters include *RGS14*, *MRPL20*, *PTK2B*, *TNFRSF4, TNFRSF18* and *CDC42SE2* up-regulation, and *CISD1*, *EDN3*, *RORC*, and *PLA2R1* down-regulation. PC1 of the entire set of 26 genes results in a TRS_UC_ that discriminates colectomy from non-progressors at p=1×10^−28^ (Fig. 3b). A score above 3.24 has a sensitivity of 90% and specificity of 95% (Fig. 3c), generating a positive predictive value of 55%, which is nine times the prevalence of the rate of progression in the study. Corresponding likelihood ratios for positive and negative prediction are 18 and 10 respectively. TRS_UC_ also performs as well as the composite PC1 of all 2,500 differentially expressed genes.

**Figure 3.**
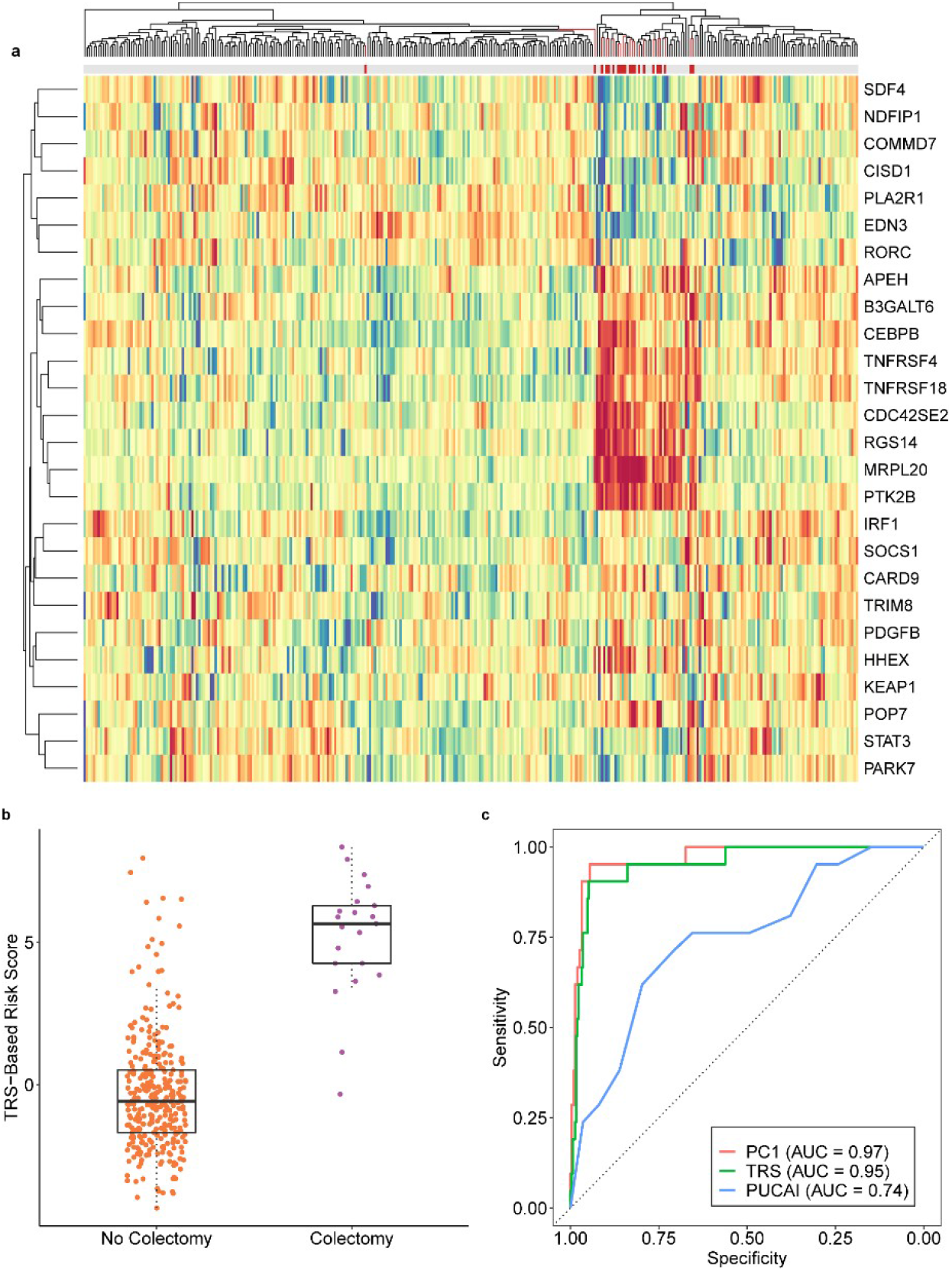
Development of a Transcriptional Risk Score for Colectomy. (a) Heatmap of baseline rectal expression of 26 genes with evidence that the GWAS peak is the same as a blood eQTL (coloc H4 > 0.8), red high expression and blue low. The gray bar at the top indicates colectomy status, highlighting a cluster of patients for whom most of the genes are differentially expressed in the cases (red bars). (b) PC1 of the genes generates a TRS that is highly discriminatory between colectomy and non-colectomy at baseline; p=1×10^−28^. Boxplots indicate 1^st^ and 3^rd^ quartile as box ends, with center median line and whiskers extending to farthest point within 1.5 times the interquartile range. (c) Receiver operating characteristic curve contrasting sensitivity and specificity for colectomy showing that both the TRS (green) and PC1 of all differentially expressed genes (red) have high accuracy (AUC > 0.95), compared with PUCAI, a commonly used clinical disease severity index.

We replicated these findings in an independent adult ulcerative colitis cohort from Mt Sinai Medical School in New York^**27,28**^. PC1 of the rectal expression of 146 genes strongly correlated with the PROTECT PC1_col_ signature highly significantly (p=0.0015) distinguished 10 patients who have had colectomy from the remaining 201 (Fig. S4a), with the majority of genes differentially expressed in the same direction. Similarly, a TRS derived from the GWAS-associated 26 transcripts showed a strong trend toward differentiation of colectomy cases in the adult cohort (Fig. S4b), which was also significant (p=0.010) after removal of two outliers characterized by aberrant expression of *CDC42SE2*, the only transcript in the list above which disagreed in direction of effect between the two studies.

Examination of the expression of colectomy-associated genes in a single cell RNAseq dataset obtained from rectal biopsies provides strong evidence that both epithelial and immune cells contribute to the risk of disease progression (Fig. S5). Most of the genes are strongly expressed in just one or two of the 22 identified cell types, seven of which are notable for an excess of colectomy associated genes: plasmocytoid dendritic cells, immunoregulatory T-cells, ILC1/3 innate immune cells, and inflammatory macrophages from the immune compartment, and fibroblasts, secretory epithelial, and endothelial cells from the gut itself. The correlated expression of these gene sets suggests that risk of colectomy may in part reflect abnormal relative abundance of these cell types. On the other hand, each of these cell types is also represented in the single cell profiles of the TRS genes, which were selected on the basis of joint eQTL and GWAS associations and hence are likely to be related to pathology through cis-regulatory effects. Prospective scRNAseq studies will likely reveal more insight into the cellular and genetic basis of the transcriptional risk of adverse disease progression.

Despite the strong contribution of trans-regulation to the TRS_UC_ score, implied by the covariance of expression of the genes, the conjunction of GWAS and eQTL signals suggests that it may be possible to also predict disease progression from genotypes alone. To evaluate this, we performed a transcriptome-wide association study^**15,16**^ using Dirichlet Process Regression (DPR) implemented in TIGAR^**29**^ to capture the effects of all polymorphisms within 1Mb of each transcript expressed in the PROTECT rectal biopsies, and then used the weights to predict gene expression in the White British subset of the UK Biobank^**21**^. We tested for differential predicted gene expression in 70% of the samples, and discovered ~800 genes either up- or down-regulated in ulcerative colitis cases relative to non-IBD controls. A predicted polygenic transcriptional risk score (PPTRS_UC_) was then derived as a weighted sum of the effect sizes of the minor alleles (which polarizes effects of alleles that increase or decrease expression in cases), and applied to the held-out 30% validation sample, as well as to the PROTECT genotypes. Figure 4a shows that the PPTRS efficiently discriminates UC cases from non-IBD controls in UK Biobank (p<10^−219^), and remarkably that it also discriminates the colectomy cases in both UK Biobank and PROTECT (p=0.002 and 0.006 respectively, p-values computed using Kruskal-Wallis test in R). That is to say, as with the observed gene expression, colectomy cases are distinguished by a trend toward yet more extreme predicted gene expression. The same trend was replicated in a larger and completely independent NIDDK-IBDGC colectomy cohort^**30,31**^, consisting of 2838 non-IBD controls, 2298 cases diagnosed as UC, and 753 known colectomy cases. The rectum-based PPTRS in this cohort discriminates UC cases from non-IBD controls (p=8.5×10^−07^) as well as UC from colectomy (p=0.0025) (Fig. 4a).

**Figure 4.**
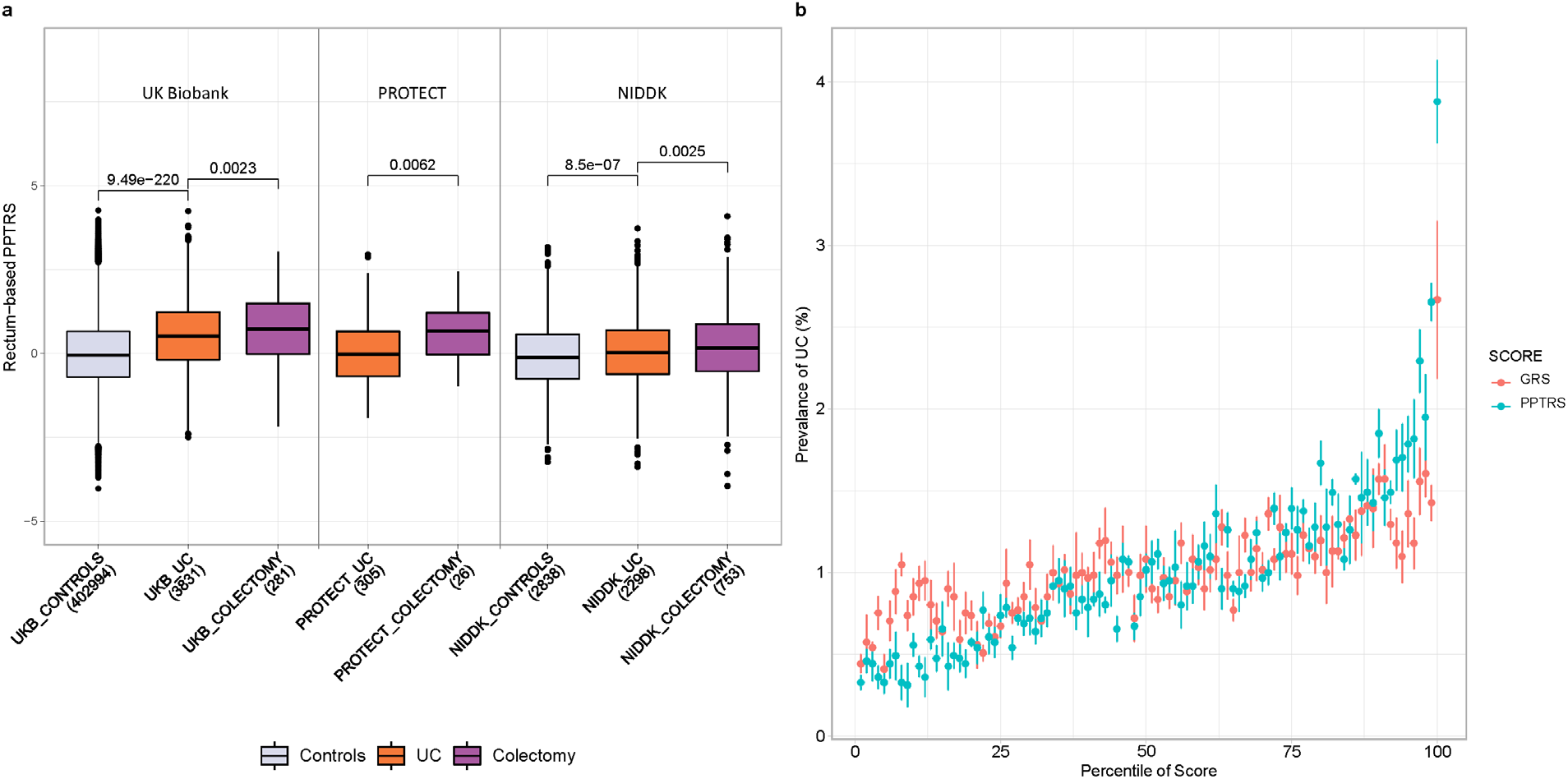
Properties of a Predicted Polygenic Transcriptional Risk Score (PPTRS). (a) PPTRS developed from predicted gene expression in PROTECT used to identify predicted differentially expressed genes in the UK Biobank. The weighted sum of 820 predicted gene expression values clearly separates controls from ulcerative colitis cases in the UK Biobank, PROTECT and NIDDK studies, while colectomy cases have even more highly elevated scores. (b) Prevalence versus Percentile plots for a Polygenic Risk Score based on 6396 genotypes for UC (red) and the PPTRS (green), showing enhanced prevalence for the upper deciles of the PPTRS. Whiskers show standard error of mean from 5-fold cross-validation.

Furthermore, PPTRS_UC_ provides enhanced discrimination of cases and controls in the UK Biobank, as shown in the prevalence vs. risk score percentile plots in Fig. 4b. Whereas the top percentile has three-fold higher prevalence than the median using a PRS with 6,396 UC SNPs from summary statistics of the European UC GWAS meta-analysis^**32**^ (pruned using PLINK at p-value < 0.001, LD *r*^2^ > 0.5), the top percentile of PPTRS_UC_ is four-fold higher, and higher prevalence is inferred for the top 20% of the entire cohort. Negative predictive values are similar for both scores.

Although colectomy status was not incorporated into either the DPR-based prediction of gene expression or the computation of PPTRS_UC_, the fact that the prediction and testing datasets are both from PROTECT could confound the interpretation with an element of circularity. We thus used the GTEx study^**33**^ transverse colon samples (n=368) to generate independent prediction models, which were then run through the same pipeline to generate a confirmatory PPTRS_UC_. Table 1 shows that this score was almost as good as the PROTECT-derived one in predicting colectomy in the UK Biobank, PROTECT and NIDDK studies (p=0.011, p=0.007 and p=0.006 respectively). Furthermore, neither cortex nor muscle-derived PPTRS from GTEx significantly predicts progression to colectomy (Table S1).

**Table 1.**
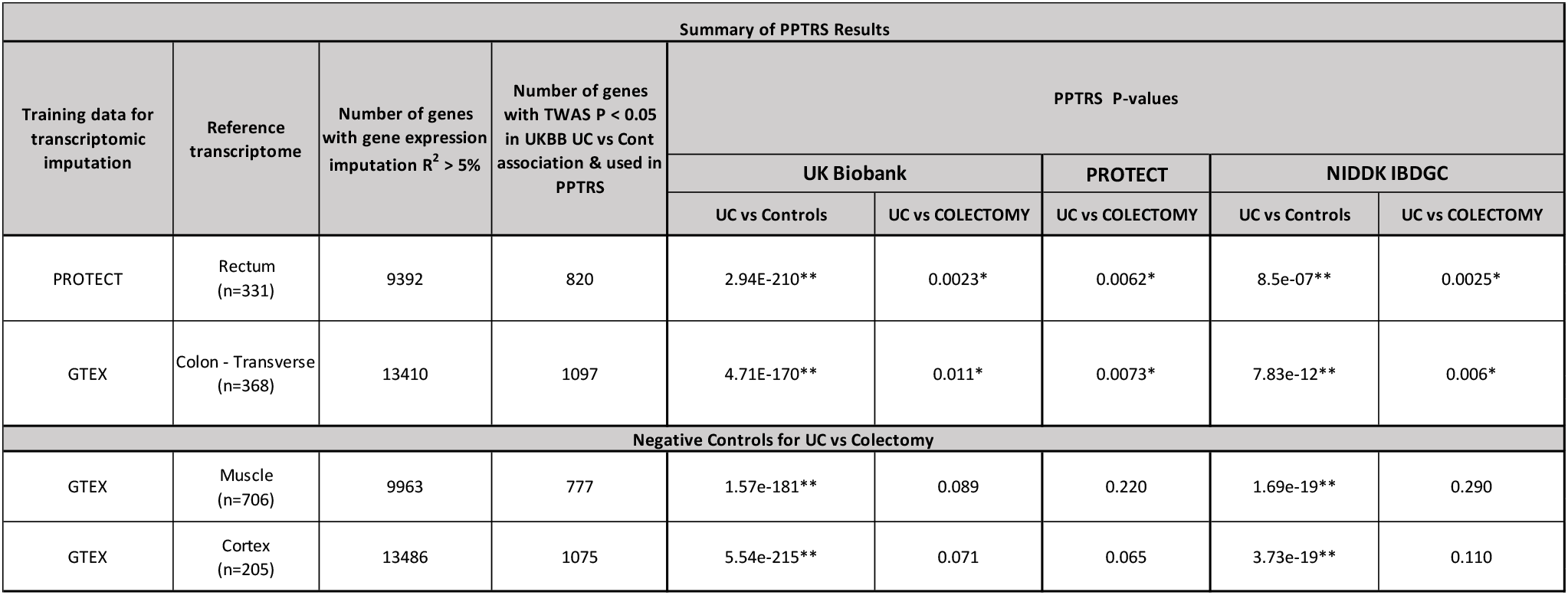
Summary of PPTRS results.

Our results highlight the potential of transcriptional profiling for prediction of colectomy in ulcerative colitis. Direct measurement of rectal biopsy RNA provides a highly discriminatory signature observed in almost all children who will need surgery, and which predicts the adverse outcome in up to half of all cases. This expression profile reverts to a healthier state regardless of immunological therapy within one year. Although much of the mis-expression is thus associated with disease status and due to trans-regulation^**34**^, we nevertheless show that prediction of gene expression from cis-linked SNPs is sufficient to generate a polygenic risk score that outperforms one based purely on GWAS associations. Our results are limited by the relatively small sample size of colectomies in the PROTECT study, which is nevertheless the largest treatment-naïve inception cohort to date. It is likely that more widespread sampling of this and other forms of inflammatory bowel disease will yield even more accurate predictors of disease progression, influencing personalized therapeutic decisions.

## METHODS

### The PROTECT cohort

428 participants aged 4 to 17 years were enrolled from 29 centers across North America into the PROTECT study upon clinical, histological, and endoscopic diagnosis of ulcerative colitis. Patients with disease extent beyond the rectum, a Pediatric Ulcerative Colitis Activity Index (PUCAI) score of ≥ 10, no prior therapy for colitis, and negative enteric bacterial stool culture were eligible to participate. All baseline assessments and sample collections were performed prior to the initiation of therapy. Initial treatment with mesalamine, oral corticosteroids, or intravenous corticosteroids was decided based on mild, moderate, or severe PUCAI. Following the baseline assessment, follow-up assessments were performed at 4, 12, and 52 weeks, with other therapeutic interventions administered based on guidelines for need additional medical therapy. Study parameters are described in further detail in Hyams et al (1).

### RNAseq data processing and differential expression analyses

RNA was isolated from 340 rectal biopsies taken at baseline and 92 rectal biopsies taken at week 52 follow-up. RNAseq was performed with the Lexogen QuantSeq 3’ platform. Using FastQC, the single end 150 bp reads were trimmed and adapters were removed (2). Reads were mapped to human genome hg19 using hisat2, and the aligned reads were converted into read counts per gene with SAMtools and HTSeq in the default union mode (3),(4),(5). The raw read counts were normalized via trimmed mean of M-values normalization with the edgeR R package (6).

Expression of the sex-specific genes RPS4Y1, EIF1AY, DDX3Y, KDM5D, and XIST was used to validate the gender of each individual, resulting in the removal of two mismatches. Further adjustment and removal of batch effects was performed with surrogate variable analysis (SVA) combined with supervised normalization (SNM) (7),(8). Race, gender, initial treatment group, time of sampling, and week 52 colectomy status were modeled with the SVA R package, where initial treatment group, time of sampling, and week 52 colectomy status were protected variables, which resulted in the identification of 28 confounding factors. Of these, five variables significantly correlated with protected variables were preserved, while the remaining 23 were statistically removed with SNM. Two individuals that were outliers in a principal component analysis of total gene expression were removed.

Differential gene expression testing was performed based on colectomy status with the voom R package. Log fold change and Benjamini-Hochberg adjusted p-values were obtained for all genes. The first principal component of the top 150 genes differentially expressed at baseline between patients who required colectomy by week 52 follow-up (n= 21) and patients who did not (n= 310) formed the gene expression-based risk score for colectomy (PC1_col_). This score is moderately correlated (r=0.46) with PC1 of overall expression of genes differentiating UC cases and controls, reported by Haberman et al (2019) (reference 7 in main text).

Cross validation for PC1_col_ was performed by randomizing colectomy status amongst individuals prior to differential gene expression testing and calculation of PC1_colRand_, as in the calculation for PC1_col_. ANOVA was performed between randomized colectomy and non-colectomy individuals, with results from 1000 such tests reported in Fig. S1.

We compared expression of the genes comprising PC1_col_ at baseline and week 52 with Mayo score as a marker for mucosal healing (Fig. S2). PC1_col_ was calculated as previously described in the subset of individuals with baseline gene expression. Additionally, a restricted PC1_col-wk52_ was calculated by finding PC1 of the 150 genes used in the calculation of PC1_col_, within the subset of individuals with week 52 gene expression. Change in PC1 score was simply calculated as the difference between PC1_col_ and PC1_col-wk52_. All p-values were generated with analysis of variance (ANOVA) tests.

Transcriptional Risk Scores (TRS), first introduced by Marigorta et al. (9) for discriminating IBD cases versus controls, capture the summation of polarized expression of genes incorporated based on both proximity to IBD GWAS hits and presence of eQTLin peripheral blood. We generated the TRS with four different strategies, all of which gave similar highly significant differentiation between colectomy and no colectomy samples. Model 1 was a GLM using the top 9 genes *RGS14, APEH, MRPL20, POP7, CDC42SE2, RORC, EDN3, PTK2B,* and *STAT3* that differentiate patients by colectomy status (*p* < 0.1), essentially the sum of the z-scores weighted by their magnitude of differential expression. Model 2 was a GLM using the 10 genes discussed in the text due to strong co-regulation and association with colectomy. Models 3 and 4 were based on all 26 genes, generated with a weighted GLM or simple PC1 score, respectively. All four scores are highly correlated, r>0.8, indicating that they are capturing similar aspects of differential expression (Fig. S7). We report Model 4 in the text. This TRS is highly correlated with PC1_col_ (r=0.64).

Relative proportions of epithelial and immune contributions to total rectal gene expression reported in Fig. S3 were evaluated by computing PC1 of the expression of 200 genes upregulated specifically in the total epithelial or immune components of the single cell gene expression dataset reported by Smillie et al (10). We checked each PC to ensure that positive values associate with elevated expression of the respective genes, and compared the values at Baseline and Week 52.

### Replication of colectomy risk score and cell-type enrichment

Surgical specimens from 210 ulcerative colitis patients undergoing bowel resection for IBD at Mount Sinai Health System and affiliated clinicians were recruited to be part of the Mount Sinai Crohn’s and Colitis Registry (MSCCR) between December, 2013 and September, 2016 as described (11–13). The protocol required written informed consent that was approved by the Icahn School of Medicine at Mount Sinai Institutional Review Board (HSM#14-00210). Patients who were enrolled in the study were asked to provide blood and/or biopsies, which were collected during a colonoscopy planned for regular care. Clinical and demographic information was obtained through a questionnaire. Patients were treated with a range or medications, including corticosteroids, infliximab, azathioprine, and mesalamine. All macroscopically moderate-to-severely inflamed tissues were confirmed as active colitis by pathology examination provided by the Mount Sinai Hospital (MSH) Pathology Department. Freshly collected representative 0.5-cm-wide tissue fragments were isolated from surgical specimen samples, flash frozen, and stored at −80 °C.

RNA was isolated from frozen tissue using Qiagen QIAsymphony RNA Kit (cat.# 931636) and samples with RIN scores >7 were retained. One microgram of total RNA depleted of ribosomal RNA using the Ribozero kit (Illumina Cat # MRZG12324) was used for the preparation of sequencing libraries using RNA Tru Seq Kits (Illumina (Cat # RS-122-2001-48). These were sequenced on the Illumina HiSeq 2500 platform using 100 bp paired end protocol. Base calling from Images and fluorescence intensities of the reads was done in situ on the HiSeq 2500 computer using Illumina software, aiming for 70,000 paired end reads per sample. Short reads were mapped to the GRCh37/hg19 assembly (UCSC Genome Browser) with 2-pasa STAR, and processed using RAPiD, which is a RNA-seq analysis framework developed and maintained by the Technology Development group at the Icahn Institute for Genomics and Multi-scale Biology. Detailed quality control metrics were generated using the RNASeQC package. Raw count data was pre-filtered to keep genes with CPM>0.5 for at least 3% of the samples. After filtering, count data was normalized via the weighted trimmed mean of M-values and further variance stabilized using a logarithmic transformation. Normalized counts were further transformed into normally distributed expression values via the voom-transformation using a model that included technical covariates (processing batch, RIN, exonic rate and ribosomal RNA rate), while accounting for the intra-patient correlation across regions.

We repeated the transcriptional risk assessment analysis in this external dataset after normalization for gender, age, exonic RNA ratio, and rRNA level expression levels, using the *prcomp* function in R with the 150 genes from the PROTECT PC1_col_, or the 26 gene TRS. The R package ggplot2 was then used to plot the distribution of PC1 for patients who did (10 patients) or did not (201 patients) have follow-up colectomies (Fig. S4). Additionally, we performed hierarchical clustering of single-cell gene expression data to identify cell types implicated by both the PC1 and TRS gene sets. Cell types enriched for PC1 genes included plasmacytoid dendritic cells, endothelial cells, group I innate lymphoid cells, fibroblasts, and macrophages.

### SNP data processing and eQTL studies

The Affymetrix UK BioBank Axiom Array was used to perform genotyping of 424 individuals across 800,000 SNPs. Imputation was performed using IMPUTE2 software (14), after which quality control performed using PLINK was used to remove SNPs not in Hardy-Weinberg equilibrium at *p* < 10^−3^, SNPs with a minor allele frequency < 1%, or a rate of missing data across individuals > 5% (15). Approximately 7 million imputed SNPs passed these thresholds and were tested in the eQTL analysis. SNPs within 250 kb of the start and stop sites of a gene were considered to be *cis* to the gene and tested for a potential eQTL association. Mapping was performed with the mixed linear modelling method in GEMMA, which tested a set of approximately 12 million SNP-gene pairs for associations at a common *p*-value threshold of 1×10^−5^ [(16)]. Two separate comparative analyses were performed, where the initial set of eQTL mapping was performed on all 330 baseline samples and 87 week 52 follow-up samples, and the secondary analysis was performed on 78 matched samples only, where the same individual was profiled at both time points. The initial full analysis yielded 91,774 significant SNP-gene associations at baseline and 19,371 associations at week 52 follow-up, and the secondary matched analysis yielded 14,272 significant unique SNP-gene associations at baseline and 12,617 significant associations at week 52 follow-up. These were further refined to 1,317, 218, 186, and 166 peak SNP to unique gene associations, respectively.

### Single cell sequence analysis of the lamina propria

For the analyses reported in Supplementary Fig. S5, we analyzed a total of 34,157 cells from paired inflamed rectum (n = 4) and uninflamed sigmoid colon (n = 5) from 4 UC patients undergoing treatment at Mount Sinai Hospital. Resected tissue biopsies were collected in ice cold RPMI 1640 (Corning Inc.) and processed within one hour after termination of the surgery. To limit biased enrichment of specific cell populations related to local variations in the intestinal micro-organization, we pooled twenty mucosal biopsies sampled all along the resected specimens using a biopsy forceps (EndoChoice). Epithelial cells were dissociated by incubating the biopsies in a dissociation medium (HBSS w/o Ca^2+^ or Mg^2+^ (Life Technologies) with HEPES 10mM (Life Technologies) and enriched with 5mM EDTA (Life Technologies)) at 37°C with 100 rpm agitation for two cycles of 15 min. After each cycle, the biopsies were vortexed vigorously for 30 seconds, and washed in complete RPMI media equilibrated at RT. They were transferred to digestion medium (HBSS with Ca^2+^ Mg^2+^, FCS 2%, DNase I 0.5mg/mL (Sigma-Aldrich) and collagenase IV 0.5mg/mL (Sigma-Aldrich)) for 40 min at 37°C with 100 rpm agitation. After digestion, the cell suspension was filtered through a 70mm cell strainer, washed in DBPS / 2% FCS / 1mM EDTA and spun down at 400 g for 10 min. After red blood cell lysis (BioLegend), dead cells were depleted using the dead cell depletion kit (Miltenyi Biotec, Germany), following manufacturer’s recommendations. Viability of the final cell suspension was calculated using a Cellometer Auto 2000 (Nexcelom Biosciences) with AO/PI dye. The exclusion was routinely 70% or higher live cell rate.

Single cells were processed through the 10X Chromium platform using the Chromium Single Cell 3′ Library and Gel Bead Kit v2 (10X Genomics, PN-120237) and the Chromium Single Cell A Chip Kit (10X Genomics, PN-120236) as per the manufacturer’s protocol. In brief, 10,000 cells from single cell suspension were added to each lane of the 10X chip. The cells were partitioned into gel beads in emulsion in the Chromium instrument, in which cell lysis and bar-coded reverse transcription of RNA occurred, followed by amplification, fragmentation and 5′ adaptor and sample index attachment. Libraries were sequenced on an Illumina NextSeq 500.

We aligned reads to the GRCh38 reference using the Cell Ranger v.2.1.0 Single-Cell Software Suite from 10X Genomics. The unfiltered raw matrices were imported into R Studio as a Seurat object (Seurat v3.0.1 (17)). Genes expressed in fewer than three cells in a sample were excluded, as were cells that expressed fewer than 500 genes and with UMI count less than 500 or greater than 60,000. We normalized by dividing the UMI count per gene by the total UMI count in the corresponding cell and log-transforming. The Seurat integrated model (17) was used to generate a combined ulcerative colitis model with cells from both inflamed and uninflamed samples retaining their group identity. We performed unsupervised clustering with shared nearest-neighbour graph-based clustering, using from 1 to 15 principal components of the highly variable genes; the resolution parameter to determine the resulting number of clusters was also tuned accordingly. Cell types were assigned using known markers previously described for Crohns’ disease (18). Visualization of relative abundance of specific genes in each cell type was performed using Seurat functions in conjunction with the ggplot2 (19).

### Gene expression imputation and prediction models

We performed a transcriptome wide association study (TWAS) for association between the imputed cis-genetic component of gene expression with UC status. PROTECT (1) was used as the prediction study with both genetic and transcriptomic data from which to estimate cis-eQTL effects, which were then used to impute gene expression in the UK Biobank validation dataset. Subsequently, these predicted gene expression models were associated with UC status in the UK Biobank, and the significant ones were combined into a weighted Predicted Polygenic Transcriptional Risk Score (PPTRS) which was itself evaluated for association with UC, and secondarily with colectomy status, in PROTECT (1).

Before building the gene expression imputation models, we ensured that the prediction and validation studies were harmonized, such that the allele frequencies are correlated, by ensuring that the genotype matrix accounts correspond to the same allele in both datasets. Gene expression imputation models were built using a non-parametric Bayesian Dirichlet process regression (DPR) method (20,21) in TIGAR, which assumes a Dirichlet process prior on the effect size variance to estimate cis-eQTL effect sizes. A linear regression model was assumed for estimating cis-eQTL effect sizes:

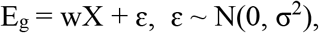

where E_g_ is the gene expression for a gene g, X is the genotype matrix for all cis-genotypes (SNPs within 1MB of the flanking 5’ and 3’ ends), w is the vector of cis-eQTL effect sizes, and ɛ is the error term assumed to be normally distributed with a mean of zero. The predicted (imputed) gene expression for gene g is computed as:

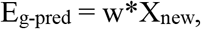

where X_new_ is the cis-genotype matrix of the new genotype data or GWAS samples and E_g-pred_ is the predicted gene expression of the new data. The imputed gene expression is the cis-genetic component of the total gene expression derived from common cis-eQTLs and does not include the trans-component, or environmental effects. TIGAR (20) has been shown to generate a 2 fold improvement in variance explained by multi-SNP models relative to just capturing the top cis-eQTLs (22), more than with similar imputation methods such as Predixcan and FUSION (23,24).

As prediction datasets, we initially utilized the PROTECT (1) cohort (rectal gene expression, n=331), confirmed with GTEX (27) transverse colon gene expression (n=368), and contrasted with GTEx muscle gene expression (n=706) and cortex gene expression (n=205) negative controls. Sigmoid colon has fewer samples, so was underpowered for these analyses, despite being closer to the rectum than transverse colon. A threshold of 5% imputation R^2^ was used to select genes with valid imputation models that were taken forward for testing in the UK Biobank and PROTECT (Fig. S6 shows boxplots of imputation R^2^ for all tissues and table S1 showing number of genes with imputation R^2^ > 5%). Note that colectomy status was not used in the modeling of either the cis gene expression, nor generation of the PPTRS, so prediction of colectomy in PROTECT from the UK Biobank score should not be circular. However, use of the GTEx colon expression to generate the imputation models ensures that prediction, validation and testing are performed with three independent datasets (GTEx, UK Biobank, and PROTECT). Further, we also replicated these results on a larger and completely independent European subset of NIDDK IBD Genetics Consortium colectomy cohort, wherein the rectum- and colon-based PPTRS discriminated UC from colectomy, while the muscle- and cortex-based PPTRS were negative controls. Finally, we also generated the PPTRS on a subset of the UK Biobank, testing it on a held-out sample with similar results.

### Transcriptome wide association study and Predicted Polygenic Risk Score (PPTRS)

For the validation dataset, the genotype data of UK Biobank was used, including 4112 Ulcerative Colitis cases and 402,994 Non-IBD Controls. The gene expression of 407,106 White British individuals was predicted using gene expression imputation models for genes with imputation R^2^ > 5%. Subsequently, a gene-based association test was performed by fitting a logistic regression model of the predicted gene expression against UC case-control status to determine the weight (log odds ratio) and p-value for each gene.

We then built a TWAS-based polygenic risk score, which we call a Predicted Polygenic Transcriptional Risk Score (PPTRS). To assess the polygenic architecture of gene expression, we adopted a TWAS threshold for differentially expressed genes with TWAS *p*-value < 0.05. The PPTRS score was constructed by computing the weighted sum of the predicted gene expression, where the weights are the log of odds ratio from TWAS of UC in UK Biobank (25). This score, as expected, highly significantly differentiates cases and controls in the UK Biobank, and surprisingly also colectomy status. The same weights were then used to generate the PPTRS in PROTECT and NIDDK cohorts, and to evaluate association with colectomy status. This procedure was repeated with the GTEx eQTL models. The contrasting polygenic risk score derived from GWAS weights, GRS_UC_, was constructed using 6,396 UC SNPs from summary statistics of the European UC GWAS meta-analysis (26) (pruned using PLINK at p-value < 0.001, LD r^2^ > 0.5 in 10kb windows with a 5-SNP sliding step).

#### NIDDK IBDGC Colectomy Cohort

Samples were genotyped on the Illumina Global Screening Array at Feinstein Institute for Medical Research (Manhasset, NY) or at the Broad Institute (Boston, MA) as a part of the National Institute of Diabetes and Digestive and Kidney Diseases Inflammatory Bowel Disease Genetics Consortium (NIDDK-IBDGC). Following stringent pre-imputation QC metrics as previously described (28), genotypes were phased using Eagle2 (29) and imputation was performed using the Michigan Imputation Server and HRC r1.1 reference panel (30, 31). Variants with estimated imputation accuracy (Rsq)<0.3 and minor allele frequency >0.1% were excluded post-imputation, leaving 21.9 million variants available for analysis. Of the total 16,024 NIDDK IBDGC samples available post-QC, 14,659 were of European ancestry (defined as EUR Admixture proportion ≥ 0.70 (32). These included 2838 non-IBD controls, 2298 UC diagnosed cases (1325 established non-colectomy), and 753 known colectomy cases. The predicted polygenic risk score for colectomy was computed on these samples using predicted gene expression from the cis-eQTL weights calculated with DPR on the rectal gene expression from PROTECT, or alternatively colon, cortex and muscle gene expression from GTEX. The TWAS weights for inclusion in the PPTRS_col_ from the UK Biobank are reported in Table S1, with code provided by S.N. to T.H.

## Ethics statement

Each site’s institutional review board approved the protocol and safety monitoring plan. Informed consent or assent was obtained for each participant.

## Data accessibility

The RNAseq data for this study has been deposited to the NCBI GEO database, series “GSE150961”. Data will be made completely openly accessible upon publication.

## Code availability statement

No custom algorithms or software were utilized for this study, but the corresponding authors will gladly share parameters used upon request. Code for computation of the PPTRS is available at the following github link: https://github.com/sn-GT/Measured-and-predicted-TRS.git.

## ACKNOWLEDGEMENTS

Support for this study was provided by NIDDK through grants U01DK095745, R01DK119991, P01 DK046763, U01 DK062413], U24DK062429, and U01DK062422, as well as the Leona M. and Harry B. Helmsley Charitable Trust. The authors thank Urko Marigorta for his counsel in development of TRS, Frank Hamilton, MD, Dana Anderson, MD, James Everhart, PhD, Jose Serrano, MD, PhD, and Stephen James, MD from NIDDK for their guidance. This research has been conducted using the UK Biobank Resource under Application Number 17984 to GG. The authors thank PROTECT site investigators for patient recruitment and data gathering, to the research coordinators at the investigative sites for their tireless attention, and to the patients and families who graciously agreed to participate.

---

**Supplementary Table 1**

Supplementary Table S1.xlsx: Summary of PPTRS results including list of genes in each tissue.

**Supplementary Table 2**

Supplementary Table S2.xlsx: Summary of GSEA pathway results.

**Supplementary Table 3**

Supplementary Table S3.xlsx: Summary of peak eQTL identified in baseline and week 52 cohorts.

**Figure S1.**
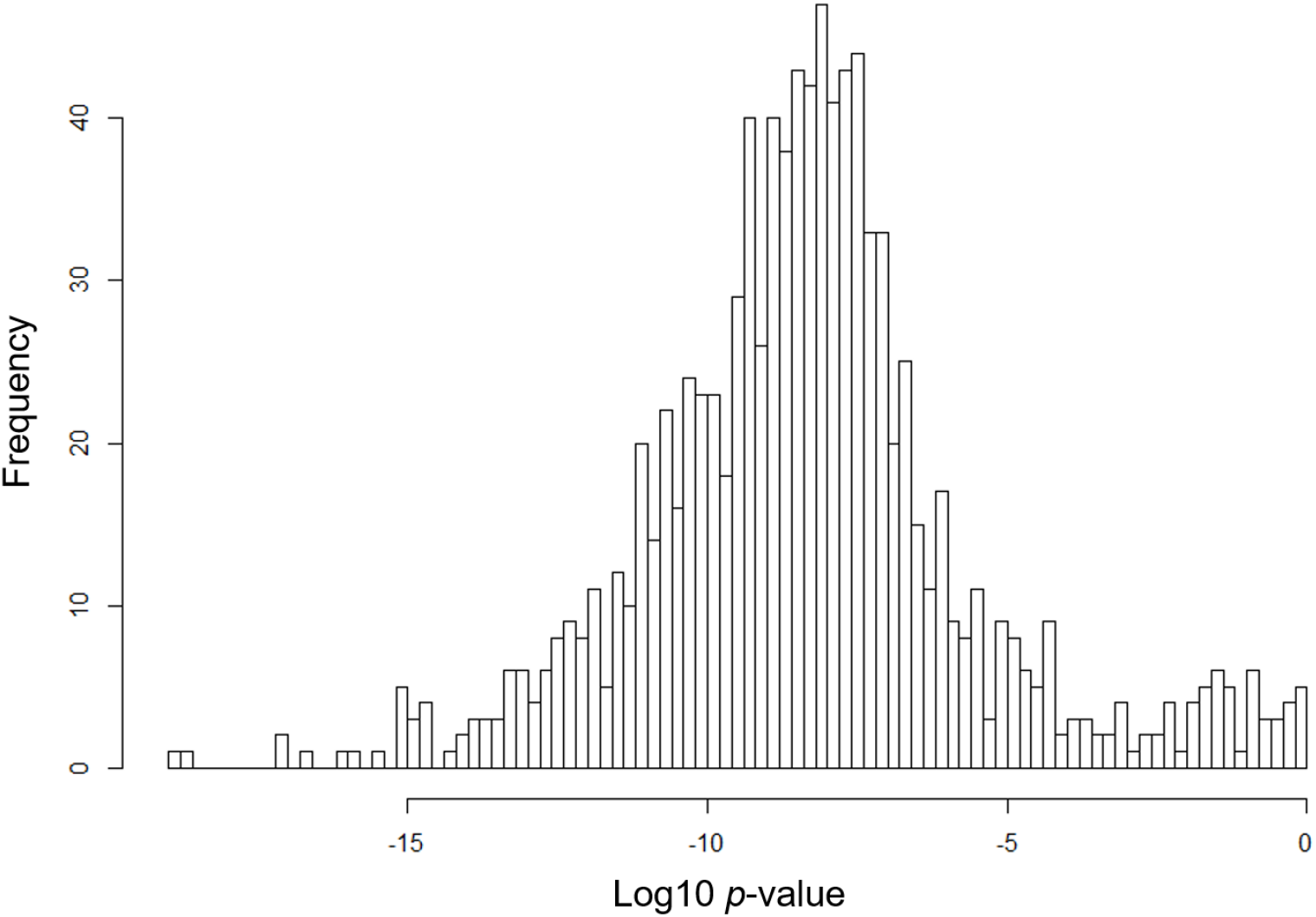
Permutation of PC1_col_. Colectomy status was randomized prior to differential expression testing and calculation of PC1_colRand_. Histogram shows frequency of log10 p-value for ANOVA test of PC1_colRand_ between randomized colectomy and non-colectomy individuals in 1000 trials. Scores tend to be significant because the PC1 is derived from transcripts that are generally significant by chance in the permuted data. However, the significance is orders of magnitude less than that derived from the actual colectomy data: PC1_col_ true p = 2×10^−45^.

**Figure S2.**
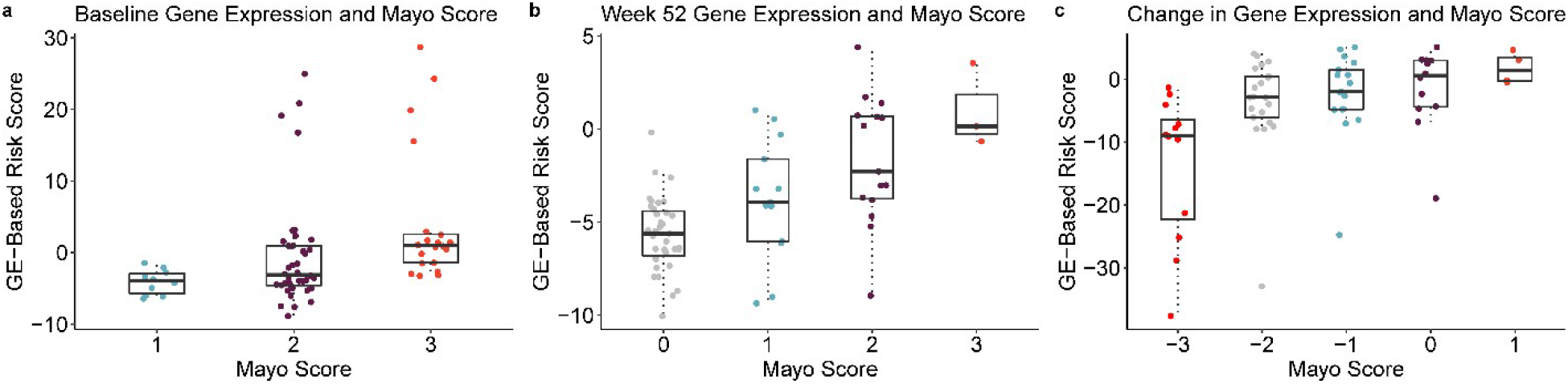
Associations between PC1_col_ and Mayo score. All boxplots indicate 1^st^ and 3^rd^ quartile as box ends, with center median line and whiskers extending to farthest point within 1.5 times the interquartile range. (a) PC1_col_ calculated on baseline gene expression with baseline Mayo score; p=0.004. (b) PC1_col_ calculated on week 52 gene expression with week 52 Mayo score; p=8.73×10^−8^. (c) Change in PC1_col_ and Mayo score from baseline to week 52; p=4×10^−4^.

**Figure S3.**
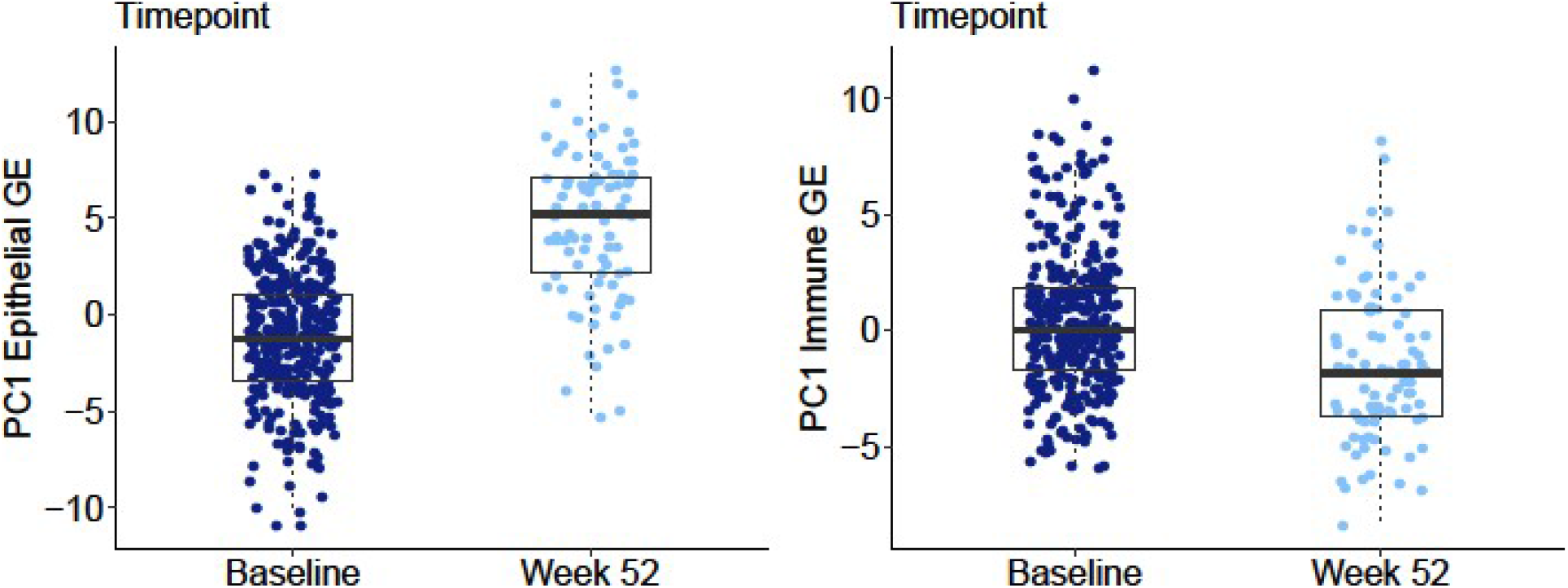
Switch in proportions of epithelial and immune components of rectal gene expression between baseline and week 52 follow-up. All boxplots indicate 1^st^ and 3^rd^ quartile as box ends, with center median line and whiskers extending to farthest point within 1.5 times the interquartile range. First principal components of 200 genes differentially expressed between the two tissue compartments in [Supplement ref. 27] were calculated and polarized such that PC1 reflects elevated expression of the genes. These results imply that immune activity is suppressed at week 52, and epithelial activity relatively elevated.

**Figure S4.**
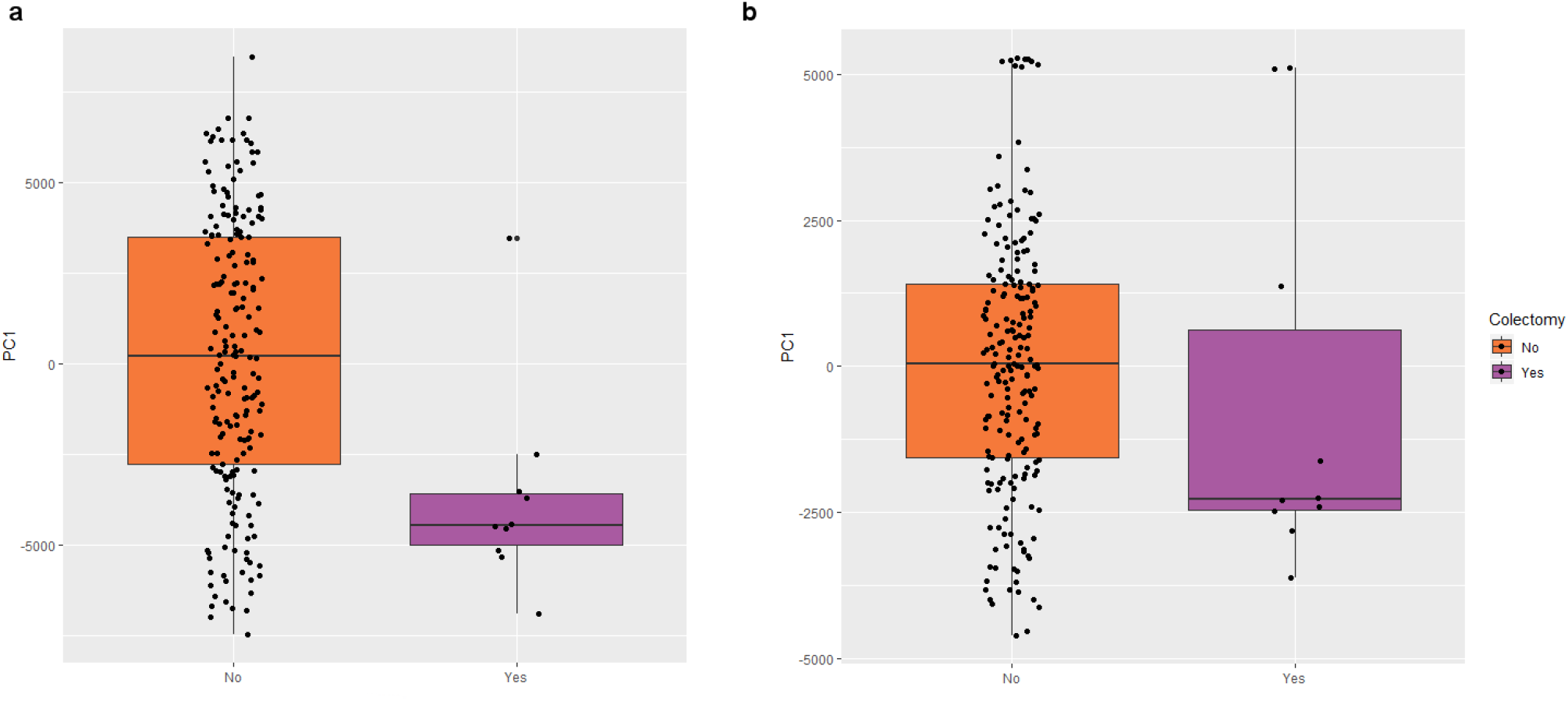
Replication of transcriptional risk prediction in the Mt Sinai cohort. All boxplots indicate 1^st^ and 3^rd^ quartile as box ends, with center median line and whiskers extending to farthest point within 1.5 times the interquartile range. (a) PC1 of colectomy-associated genes in Mt Sinai significantly differentiates colectomy (purple) from non-colectomy (orange). (b) TRS_UC_ developed from IBD GWAS-associated genes also predicts progression to colectomy in the Mt Sinai cohort. Two outlier samples reduce the significance, which is p=0.01 for the remaining samples.

**Figure S5.**
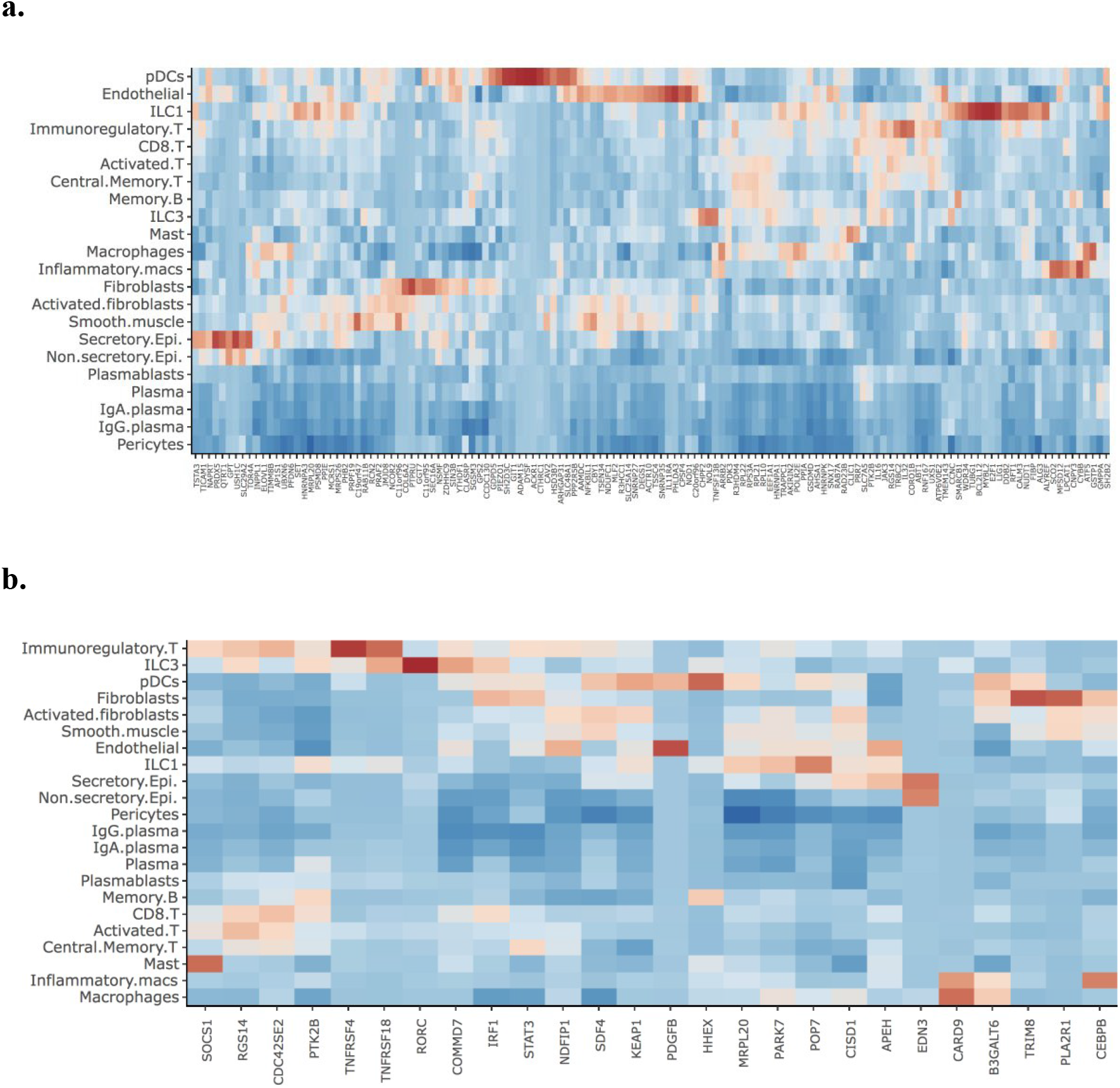
Cell-type specific expression of colectomy-associated genes. (a) Heat map showing up-regulation (red) of each gene contributing to PC1 in a rectal scRNAseq dataset. Dozens of genes are enriched in seven cell-types. (b) Similar analysis but for the TRS_UC_ genes. Note the similarity of the cell-types showing enrichment, and the absence of B-cell or plasma cell signals in both.

**Figure S6.**
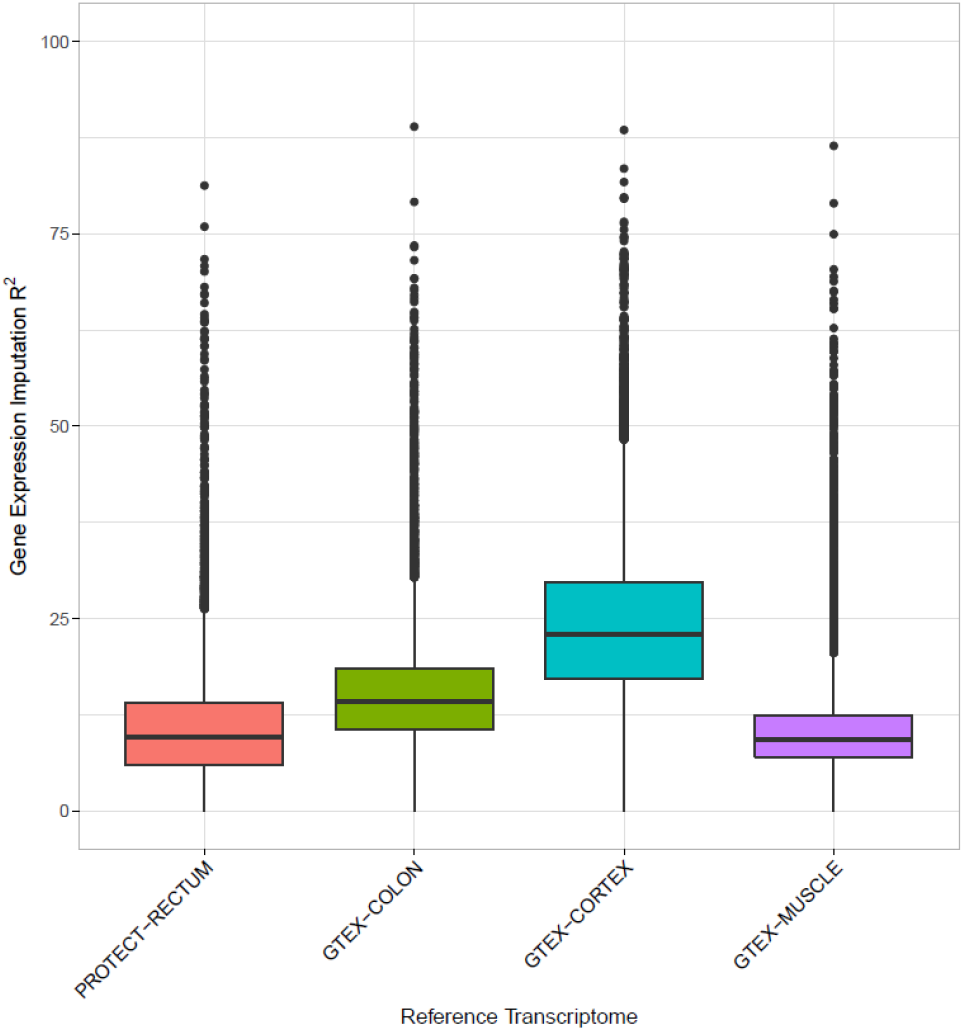
Distribution of R^2^ values for gene expression prediction models from each tissue. Each boxplot shows the median value of the variance in gene expression explained by DPR prediction with upper and lower hinge representing first and third quartiles (25^th^ and 75^th^ percentiles). The upper and lower whiskers extends no further than 1.5 × IQR (inter-quartile range) and data points beyond the end of whiskers are outliers.

**Figure S7.**
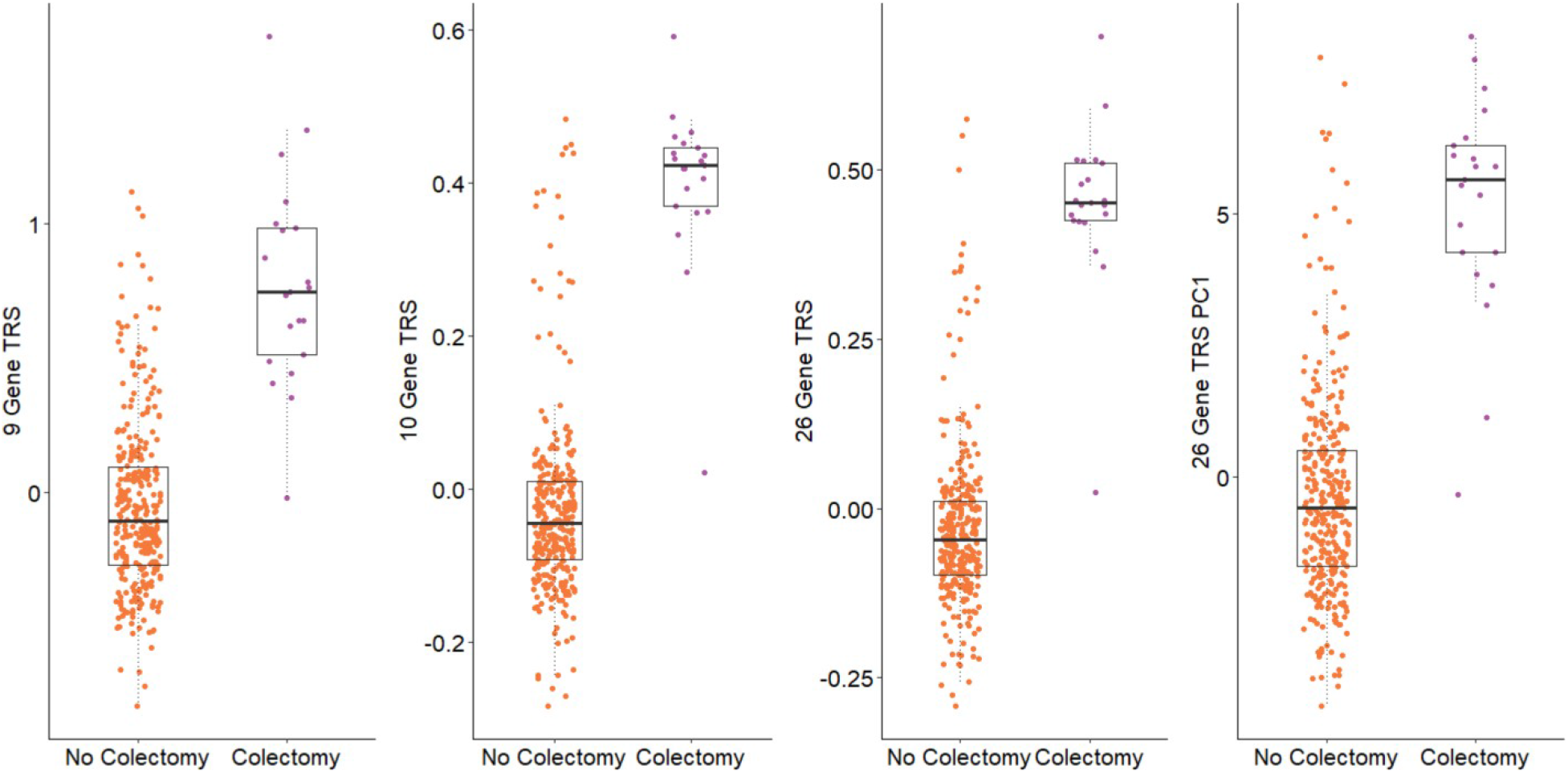
Comparison of TRS generated with different subsets of genes. Each plot shows the computed TRS for each individual who did or did not require colectomy during the study period. All boxplots indicate 1^st^ and 3^rd^ quartile as box ends, with center median line and whiskers extending to farthest point within 1.5 times the interquartile range. (a) 9 gene TRS for genes significantly differentiated by status at p<0.1; p=2×10^−25^. (b) 10 gene TRS for genes highlighted in the text as the major clusters of up- and down-regulated in colectomy; p=8×10^−43^. (c) 26 gene TRS as sum of z-scores weighted by the magnitude of differential expression; p=9×10^−49^. (d) TRS computed simply as PC1 of the 26 genes; p=1×10^−28^.

## Notes

https://www.ncbi.nlm.nih.gov/geo/query/acc.cgi?acc=GSE150961

### Competing Interest Statement

JSH has served on an advisory board for Janssen and is acting as a consultant for AbbVie, Takeda, Lilly, Boehringer-Ingelheim, Allergan, Pfizer, Receptos, and AstraZeneca. SDT has been a member of an independent data monitoring committee for Lycera Corporation. AMG has received research support from AbbVie; been a consultant for AbbVie, Celgene, Janssen, Lilly, Pfizer, and Takeda; and been a speaker for AbbVie, Janssen, and Shire. NSL has been a consultant for AbbVie. CGS has been a consultant for AbbVie. JM has been a consultant for Janssen, Celgene, and Lilly. JRR has been a consultant for AbbVie, Celgene, Janssen, Luitpold, and Pfizer, and received grant funding from Janssen and AbbVie. ASP has participated in speakers bureaus AbbVie and Janssen. MBH has received research grants from Genentech, AbbVie, Shire, Takeda, Mallinkrodt, Janssen, and Gilead. PAR has been a consultant for Shire and Leutpold; been a speaker for AbbVie; and received research support from TechLab. SK has been a consultant for Janssen and UCB. LAD has received grant support from AbbVie and Janssen. DPBM and TH are faculty members at Cedars-Sinai Medical Center. EM is an employees at Cedars-Sinai. Cedars-Sinai has financial interests in Prometheus Biosciences, Inc., a company which has access to the data and specimens in Cedars-Sinai MIRIAD Biobank. Prometheus Biosciences, Inc. seeks to develop commercial products. DM is a paid consultant and shareholder of Prometheus Biosciences, Inc. DM has consulted for Pfizer, Gilead, Palatin Technologies, Bridge Biotherapeutics, and Takeda. All other authors declare no competing interests.

## REFERENCES

Alexander, D.H., Novembre, J., Lange, K. Fast model-based estimation of ancestry in unrelated individuals. Genome Res. 19: 1655–1664 (2009).

Anders, S., Pyl, P.T., & Huber, W. HTSeq--a Python framework to work with high-throughput sequencing data. Bioinformatics 31, 166–169 (2015).

Andrews S. FastQC: a quality control tool for high throughput sequence data. https://www.bioinformatics.babraham.ac.uk/projects/fastqc/.(2010)

Aragam, K.G., Dobbyn, A., Judy, R., Chaffin, M., Chaudhary, K., Hindy, G., et al. Limitations of contemporary guidelines for managing patients at high genetic risk of coronary artery disease. J Am Coll Cardiol. 75, 2769–2780 (2020).

Bycroft, C. et al. The UK Biobank resource with deep phenotyping and genomic data. Nature 562, 203–209 (2018).

Damask, A., Steg, P.G., Schwartz, G.G., Szarek, M., Hagström, E., et al; Regeneron Genetics Center and the ODYSSEY OUTCOMES Investigators. Patients with high genome-wide polygenic risk scores for coronary artery disease may receive greater clinical benefit from alirocumab treatment in the ODYSSEY OUTCOMES Trial. Circulation 141, 624–636 (2020).

Das, S., et al. Next-generation genotype imputation service and methods. Nat Genet. 48: 1284–1287. (2016).

Gamazon, E.R., et al. A gene-based association method for mapping traits using reference transcriptome data. Nat Genet. 47, 1091–1098 (2015).

Giambartolomei, C. et al. A Bayesian framework for multiple trait colocalization from summary association statistics. Bioinformatics 34, 2538–2545 (2018).

Graham, D.B., Xavier, R.J. Pathway paradigms revealed from the genetics of inflammatory bowel disease. Nature 578, 527–539 (2020).

GTEx Consortium. Genetic effects on gene expression across human tissues. Nature 550, 204–213 (2017).

Gusev, A. et al. Integrative approaches for large-scale transcriptome-wide association studies. Nat. Genet 48, 245–252 (2016).

Gusev, A. et al. Transcriptome-wide association study of schizophrenia and chromatin activity yields mechanistic disease insights. Nat Genet 50, 538–548 (2018).

Gibson, G. On the utilization of polygenic risk scores for therapeutic targeting. PLoS Genet. 15, e1008060 (2019).

Haberman, Y. et al. Ulcerative colitis mucosal transcriptomes reveal mitochondriopathy and personalized mechanisms underlying disease severity and treatment response. Nat Commun. 10, 38 (2019).

Haritunians, T., et al. Genetic predictors of medically refractory ulcerative colitis. Inflamm Bowel Dis. 16, 1830–1840 (2010).

Howie, B.N., Donnelly, P., & Marchini, J. A flexible and accurate genotype imputation method for the next generation of genome-wide association studies. PLoS Genet. 5, e1000529 (2009).

Hyams, J.S., et al. Factors associated with early outcomes following standardised therapy in children with ulcerative colitis (PROTECT): a multicentre inception cohort study. Lancet Gastroenterol Hepatol. 2, 855–868 (2017).

Hyams, J.S. et al. Clinical and biological predictors of response to standardised paediatric colitis therapy (PROTECT): a multicentre inception cohort study. Lancet 393, 1708–1720 (2019).

Kim, D., Langmead, B., Salzberg, S.L. HISAT: a fast spliced aligner with low memory requirements. Nat Methods. 12, 357–360 (2015).

Kugathasan, S. et al. Prediction of complicated disease course for children newly diagnosed with Crohn’s disease: a multicentre inception cohort study. Lancet 389, 1710–1718 (2017).

Lambert, S.A., Abraham, G., & Inouye, M. Towards clinical utility of polygenic risk scores. Hum Mol Genet. 28(R2), R133–R142 (2019).

Lee, J.C., Biasci, D., Roberts, R., Gearry, R.B., Mansfield, J.C., et al. Genome-wide association study identifies distinct genetic contributions to prognosis and susceptibility in Crohn’s disease. Nat Genet. 49, 262–268 (2017).

Leek, J.T., Johnson, W.E., Parker, H.S., Jaffe, A.E., & Storey, J.D. The sva package for removing batch effects and other unwanted variation in high-throughput experiments. Bioinformatics 28, 882–883 (2012).

Leijonmarck, C.E., Persson, P.G. & Hellers, G. Factors affecting colectomy rate in ulcerative colitis: an epidemiologic study. Gut 31, 329–333 (1990).

Lewis, C.M. & Vassos, E. Polygenic risk scores: from research tools to clinical instruments. Genome Med. 12, 44 (2020).

Li, H., et al. The Sequence Alignment/Map format and SAMtools. Bioinformatics 25, 2078–9 (2009).

Liu, J.Z. et al. Association analysis identify 38 susceptibility loci for inflammatory bowel disease and highlight shared genetic risk across populations. Nat Genet 47, 979–986 (2015).

Lloyd-Jones, L.R. et al. The genetic architecture of gene expression in peripheral blood. Am J Hum Genet. 100, 228–237 (2017).

Loh, P-R., et al. Reference-based phasing using the Haplotype Reference Consortium panel. Nat Genet. 48: 1443–1448 (2016).

McCarthy, S., et al. A reference panel of 64,976 haplotypes for genotype imputation. Nat Genet. 48: 1279–1283 (2016).

Marigorta, U.M. et al. Transcriptional risk scores link GWAS to eQTLs and predict complications in Crohn’s disease. Nat Genet. 49, 1517–1521 (2017).

Martin, J.C., Chang, C., Boschetti, G., Ungaro, R., Giri, M., et al. Single-cell analysis of Crohn’s disease lesions identifies a pathogenic cellular module associated with resistance to anti-TNF therapy. Cell 178, 1493–1508.e20 (2019).

Mecham, B.H., Nelson, P.S., & Storey, J.D. Supervised normalization of microarrays. Bioinformatics 26, 1308–1315 (2010).

Nagpal, S. et al. TIGAR: An improved Bayesian tool for transcriptomic data imputation enhances gene mapping of complex traits. Am J Hum Genet. 105, 258–266 (2019).

Naito, T., Botwin, G.J., Haritunians, T., Li, D., Yang, S., Khrom, M., Braun, J., NIDDK IBD Genetics Consortium, Abbou, L., Mengesha, E., Stevens, C., Masamune, A., Daly, M., McGovern, D.P.B. Prevalence and effect of genetic risk of thromboembolic disease in inflammatory bowel disease. Gastroenterology in press: S0016-5085(20)35276-8 (2020).

Natarajan P, Young R, Stitziel NO, Padmanabhan S, Baber U, Mehran R, et al. Polygenic risk score identifies subgroup with higher burden of atherosclerosis and greater relative benefit from statin therapy in the primary prevention setting. Circulation 135, 2091–2101 (2017).

Ndungu, A., Payne, A., Torres, J.M., van de Bunt, M. & McCarthy, M.I. A Multi-tissue Transcriptome analysis of human metabolites guides interpretability of associations based on multi-SNP models for gene expression. Am J Hum Genet 106, 188–201 (2020).

Parikh, K., Antanaviciute, A., Fawkner-Corbett, D., Jagielowicz, M., Aulicino A., et al. Colonic epithelial cell diversity in health and inflammatory bowel disease. Nature 567, 49–55 (2019).

Peters, L.A. et al. A functional genomics predictive network model identifies regulators of inflammatory bowel disease. Nat Genet 49, 1437–1449 (2017).

Purcell, S., et al. PLINK: a tool set for whole-genome association and population-based linkage analyses. Am J Hum Genet. 81, 559–575 (2007).

Robinson, M.D., McCarthy, D.J., & Smyth, G.K. edgeR: a Bioconductor package for differential expression analysis of digital gene expression data. Bioinformatics 26, 139–140 (2010).

Sandborn, W.J. et al. Colectomy rate comparison after treatment of ulcerative colitis with placebo or Infliximab. Gastroenterology 137, 1250–1260 (2009).

Schroeder, K.W., Tremaine, W.J. & Ilstrup, D.M. Coated oral 5-aminosalicylic acid therapy for mildly to moderately active ulcerative colitis. A randomized study. N Engl J Med 317, 1625–1629 (1987).

Stuart, T, et al. Comprehensive integration of single-cell data. Cell 177, 1888–1902.e21 (2019).

Suarez-Farinas, M., et al. Disease demarcation in ulcerative cohotis is associated with different patterns of gene expression. J Crohn’s Colitis 12(Suppl 1), DOP012 (2018).

Suárez-Fariñas, M., et al. Intestinal inflammation modulates the expression of *ACE2* and *TMPRSS2* and potentially overlaps with the pathogenesis of SARS-CoV-2 related disease. bioRχiv doi: https://doi.org/10.1101/2020.05.21.109124. *Gastroenterology*, in press. (2020)

Smillie, C.S., Biton, M., Ordovas-Montanes, J., Sullivan, K.M., Burgin, G., et al. Intra- and inter-cellular rewiring of the human colon during ulcerative colitis. Cell 178, 714–730.e22 (2019).

Subramanian, A. et al. Gene set enrichment analysis: a knowledge-based approach for interpreting genome-wide expression profiles. Proc Natl Acad Sci (USA) 102, 15545–15550 (2005).

Turner, D. et al. Appraisal of the pediatric ulcerative colitis activity index (PUCAI). Inflamm Bowel Dis. 15, 1218–1223 (2009).

Ungaro, R., Mehandru, S., Allen, P.B., Peyrin-Biroulet, L. & and Colombel, J-F. Ulcerative colitis. Lancet 389, 1756–1770 (2017).

Uzzan, M., et al. Mapping of B cell landscape in ulcerative colitis lesions reveals a pathogenic response that associates with treatment resistance and disease complications. Nat. Medicine 2020; Under second revision.

Wainberg, W. et al. Opportunities and challenges for transcriptome-wide association studies. Nat Genet. 51, 512–599 (2019).

Wickham, H. ggplot2: elegant graphics for data analysis. 2nd ed. Cham: Springer (2016)

Zeng, P. & Zhou, X. Non-parametric genetic prediction of complex traits with latent Dirichlet process regression models. Nat Commun 8, 456 (2017).

Zhou, X., & Stephens, M. Genome-wide efficient mixed-model analysis for association studies. Nat Genet. 44, 821–824 (2012).

